# How to read a baby’s mind: Re-imagining fMRI for awake, behaving infants

**DOI:** 10.1101/2020.02.11.944868

**Authors:** C. T. Ellis, L. J. Skalaban, T. S. Yates, V. R. Bejjanki, N. I. Córdova, N. B. Turk-Browne

## Abstract

Thousands of functional magnetic resonance imaging (fMRI) studies have provided important insight into the human brain. However, only a handful of these studies tested infants while they were awake, because of the significant and unique methodological challenges involved. We report our efforts over the past five years to address these challenges, with the goal of creating methods for infant fMRI that can reveal the inner workings of the developing, preverbal mind. We use these methods to collect and analyze two fMRI datasets obtained from infants during cognitive tasks, released publicly with this paper. In these datasets, we explore data quantity and quality, task-evoked activity, and preprocessing decisions to derive and evaluate recommendations for infant fMRI. We disseminate these methods by sharing two software packages that integrate infant-friendly cognitive tasks and behavioral monitoring with fMRI acquisition and analysis. These resources make fMRI a feasible and accessible technique for cognitive neuroscience in human infants.

---

Infants may hold the key to understanding the origins and functions of the human mind, and yet they are difficult to study because they cannot communicate verbally and have a limited repertoire of actions. Research on infant cognition has navigated these constraints by relying on simple behavioral measures such as looking^1^ and reaching^2^. This approach has been supplemented by neural measures from electroencephalography^3^ and functional near-infrared spectroscopy^4^, which provide a window into infant cognition through the scalp without requiring external behavior. Functional magnetic resonance imaging (fMRI) stands to complement these scalp-based techniques, with sensitivity throughout the whole brain (including ventral surfaces and deep brain structures) and relatively high spatial resolution that can be linked to detailed anatomy. These advantages could dramatically improve our understanding of infant cognition^5,6^, and yet fMRI has rarely been used for this purpose.

There are many challenges to collecting fMRI data from infants: head and body motion, limited attention span and fussiness, inability to understand or follow instructions, high acoustic noise levels, and a lack of analysis approaches optimized for the infant brain. Several labs have performed fMRI experiments with infants who are sedated^7^ or sleeping^8–10^. However, there have been only a handful of successes when infants are awake and behaving during cognitive tasks^11–13^. The challenges above severely limit the amount of fMRI data that can be obtained. Under the best of circumstances, it is not unusual for infants to “fuss out” after a few minutes of a task, compared to 60–90 minutes of fMRI data typically acquired from adults. In those precious few minutes, motion of the head or body, startle responses to noise, separation from parents, crying from hunger, teething or soiled diapers, and fatigue and napping, can reduce the amount of usable data to near zero. This assumes that data collection begins in the first place — some infants are not willing to lie down or wait through initial steps for the task to start. In sum, the rates of data exclusion and subject attrition seem to ensure that insufficient data are collected in order to detect what are, even in adults, relatively small and noisy signatures of cognition in the brain.

Over the past five years, we have developed a new approach for awake infant fMRI. This was made possible by innovations in various domains. First, we incorporated advances in the preprocessing and analysis of *adult* fMRI data. For example, beyond localizing functions in the brain, multivariate methods from machine learning have made it possible to extract and interpret the mental contents of neural activity patterns (e.g., specific percepts^14^, memories^15^, and decisions^16^). Second, we built on methodological and theoretical groundwork in developmental psychology. This includes infant-friendly behavioral tasks whose underlying neural foundations are mostly unknown, but for which such data could help resolve ongoing debates. Third, we combined commercially available MRI and auxiliary equipment with open-source software to create custom solutions for data acquisition, stimulus presentation, and behavioral monitoring. This allowed us to redesign the scanning environment and experimental protocols for infants from the ground up.

Some of our procedures were inspired by previous infant fMRI studies, others by studies in populations with related constraints (e.g., patients, animals), and yet others are novel and the result of extensive trial and error. No one innovation is sufficient in our experience, so here we report a family of methods that collectively allow for the robust collection of fMRI data from awake, behaving infants. In addition to describing these methods in detail, all of the code needed to implement them has been made publicly available. This includes an experiment menu system that flexibly incorporates infant-friendly cognitive tasks and seamlessly coordinates stimulus presentation, behavioral monitoring, and scanner synchronization, as well as a semi-automated pipeline tailored for the analysis of the resulting infant data. We have deployed these methods at three scanning sites, with findings from the first two completed cohorts reported here. These two fMRI datasets from awake infants have also been made publicly available. Our hope is that these software and data resources, combined with included recommendations about recruitment, safety, equipment, task design, personnel, preprocessing, and more, will help make fMRI a more feasible and prevalent technique for studying the early developing mind.

## Results

Below we emphasize two key components of the infant fMRI methods we have developed: the apparatus and procedures used in *data acquisition* (Figure 1), and the algorithms and signal processing used in *data analysis* (Figure S1). We provide four metrics to evaluate success: (1) the quantity of infant data and quality relative to adult data; (2) the reliability of BOLD activity evoked in visual tasks; (3) how these visual responses vary across preprocessing decisions in one dataset; and (4) a replication in a second dataset from a different age range and new site. Our hope is that the methods described herein, and the code to implement them we are releasing with this paper, will accelerate the adoption and refinement of awake infant fMRI.

**Figure 1:**
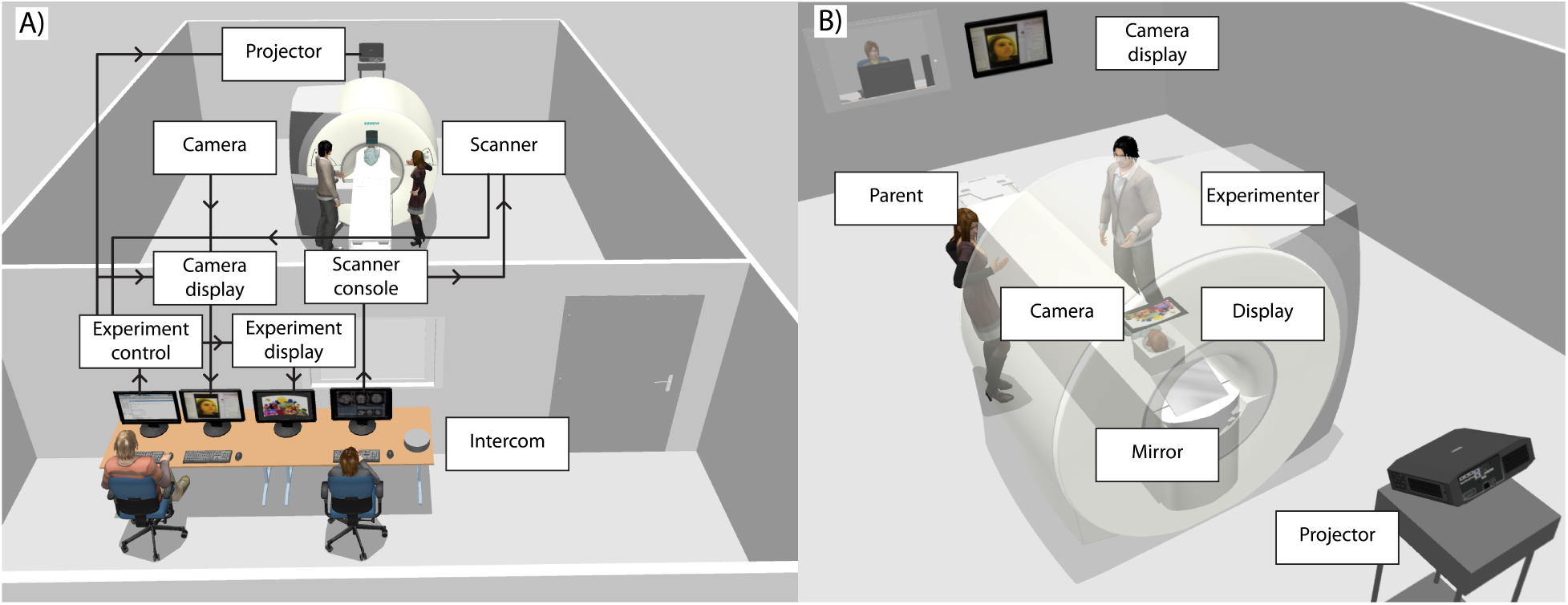
Schematic of the scanning environment. A) Overview of the setup and wiring diagram of the equipment and communications. B) Key elements inside the scanner room. One experimenter remains in the scanner room during the scan and communicates with the parent, directly monitors the infant, and coordinates with experimenters in the control room. A camera mounted in the scanner bore and a projector positioned behind and above the scanner are connected to experiment computers in the control room. The projector transmits an image down onto a mirror lying flat on the tracks of the scanner table, which reflects the image up onto the scanner bore ceiling above the infant’s face (corrected for elliptical and keystone distortions). After preparing the infant with redundant hearing protection, the experimenter places the infant’s body on a vacuum pillow on the scanner table and gently lays the infant’s head on a foam pad in the bottom of the head coil. Two other experimenters remain in the control room: one operates the experiment computers, determines which tasks to run and when, monitors the camera feed for attention and compliance, and operates the vacuum pump (not depicted); the other operates the scanner console computer, registers the participant, chooses sequences and slice alignment, and communicates with the experimenter in the scanner room.

### Data quantity and quality

We collected two types of scans in each session: an anatomical image using a T1-weighted pointwise encoding time reduction with radial acquisition (PETRA) sequence^17^, and functional data using an echo planar imaging (EPI) sequence while infants performed cognitive tasks. Although we used a standard EPI protocol, the provided experiment and analysis code is fully compatible with a range of sequence parameters, including higher resolution, faster sampling, and/or greater acceleration. Only the TR duration needs to be specified in the pipeline.

To minimize motion during anatomical scans, the child watched one or more television shows (e.g., Daniel Tiger, Sesame Street), animations (e.g., dynamic textures and shapes), and/or other videos considered engaging from a library in the experiment code. On occasion, we collected an anatomical scan when the child fell asleep during a functional scan. The PETRA sequence (part of the Siemens Quiet Suite) was more robust to motion than other anatomical sequences (e.g., MPRAGE) in early piloting, we suspect because it samples K-space radially and is short. That said, any other T1- or T2-weighted anatomical sequence could easily be integrated into the analysis pipeline by specifying the name of the scan.

Functional data were deemed usable when time-point to time-point translational motion was below a threshold of 3mm (voxel resolution). Epochs of data during a task (*blocks* in a block design or *trials* in an event-related design) were excluded if the infant’s eyes were off screen or closed more than 50% of the epoch. Eye position was determined by manual gaze coding of a video recording of the infant’s face. Note that the provided software is compatible with different video recording systems, including commercial eye-tracking software and cameras. Additional epochs were excluded at later stages of analysis when insufficient data had been collected in a task or to counterbalance conditions or timing. We occasionally kept functional scans running when the infant fell asleep during a task, though we removed the stimuli from the screen in such situations and labelled these epochs as resting data.

We report data from two cohorts of infants acquired sequentially from different sites (Figure 2). Cohort I was collected on a 3T Siemens Skyra MRI at Princeton University over a broad age range (6–36 months). Cohort II was collected on a 3T Siemens Prisma MRI at Yale University from a younger and narrower age range (3–12 months). Given the small number of studies of this type, we treated Cohort I as an exploratory sample in which to search for good analysis parameters and then applied these parameters in a principled way to Cohort II. The data were analyzed with a custom software pipeline (Figure S1), using code that we are releasing with this paper.

**Figure 2:**
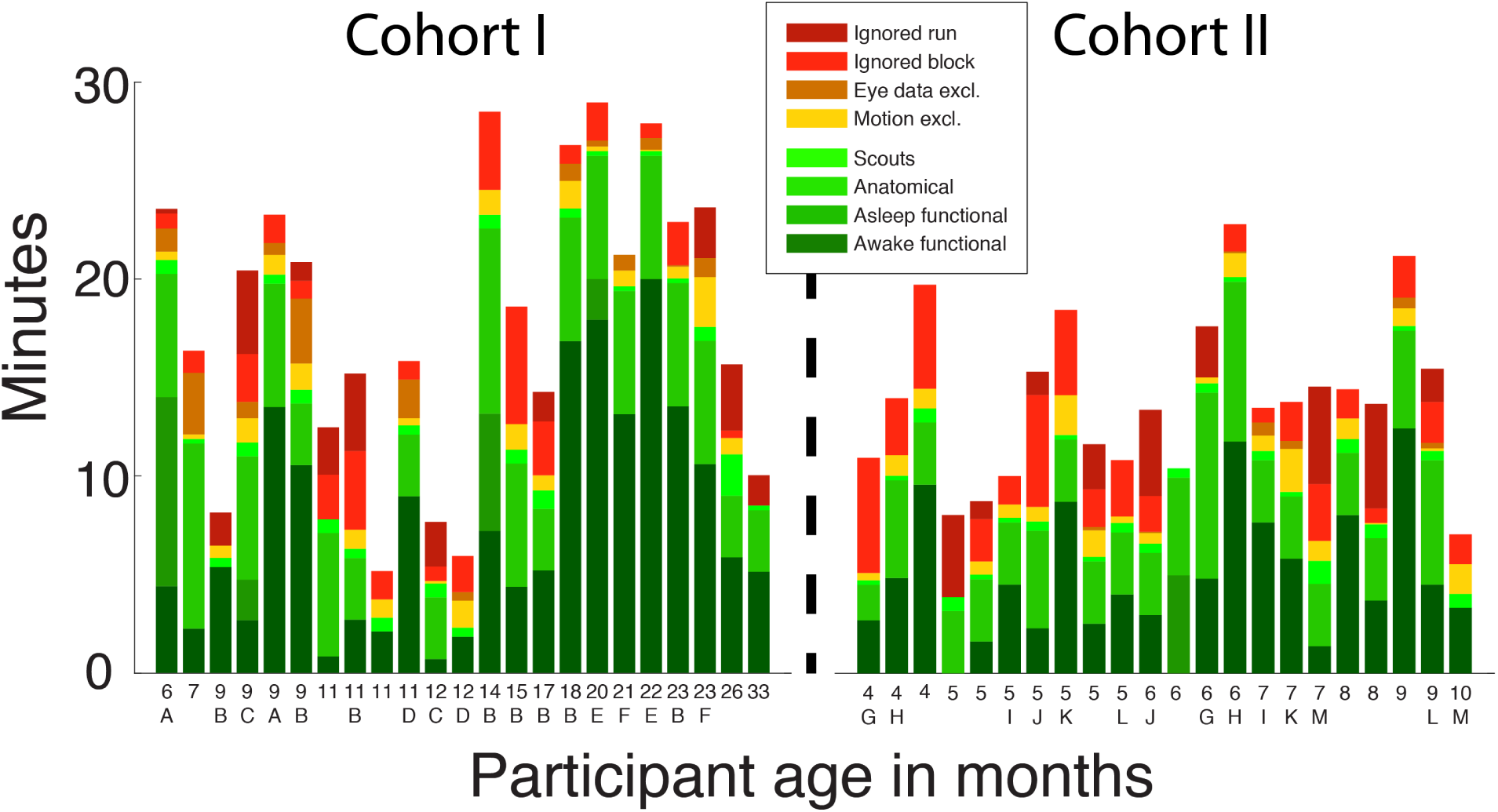
Cumulative bar plot of data retention for individual scanning sessions. The age of the child in months is shown on the *x* axis, the duration of the session on the *y* axis, including a breakdown by color for different categories of data. Individual infants who completed more than one session were assigned a letter code, which is shown beneath the age of each of their sessions (no letter means the infant from that session only participated once).

In Cohort I, we scanned 11 distinct individuals between 6 and 33 months (*M*=15.2 [*SD*=6.9]) across 23 sessions (1–8 sessions per participant). Five additional sessions could not be used because the child did not go in the scanner. Of the participants who were scanned, an average of 31.3% (13.0 mins) of the total time when experiment code was running (proxy for how long we were trying to scan) produced usable functional and/or anatomical data. After preprocessing, 16 of the 28 sessions resulted in the acquisition of at least one full experiment’s worth of usable data. Because some participants completed more than one experiment per session (range: 0–3), our overall retention rate of usable experiments per scheduled session was 1.18. Although the amount of data in minutes per experiment is low relative to adults, we obtained slightly more than one useful experiment dataset per session on average.

Beyond obtaining a reasonable quantity of data in Cohort I, we also explored whether the data were of sufficient quality. One unique aspect of our apparatus is that we did not use the top of the head coil, as typical with adults. This reduces the number of coil elements, which in principle reduces the amount of signal received from brain regions that would have been nearby. This was a deliberate decision made to increase infant comfort, provide parents an unobstructed view of the infant’s face, and enable ceiling-based visual stimulation and reliable camera-based eye-tracking — all essential in our experience. Signal loss would only be expected in the anterior portion of the brain most distant from the remaining bottom coil elements. Such loss may be partly mitigated by the small size of the infant brain, which reduces this distance. In fact, if the top coil had been used, it would have been far from the infant’s forehead, minimizing the impact of these elements on image acquisition. Other groups have addressed this problem with a custom head coil^12^. Indeed, the provided software is compatible with other visual stimulation options, including typical rear projection systems. Tools are included for calibrating the display, equating stimulus sizes across display formats, and synchronizing display timing with the video feed used for eye-tracking.

Given our non-traditional approach, we first evaluated signal quality when using only the bottom coil. A common metric for describing the quality of fMRI data is the signal-to-fluctuation-noise ratio (SFNR)^18^: the magnitude of the BOLD signal in a brain voxel relative to its variability over time. We compared voxelwise SFNR in our infant data with only the bottom coil connected against a gold standard of adult data with both bottom and top coils. The adult data were obtained from a published study^19^ using the same scanner and EPI sequence as the infants in Cohort I. We analyzed data from 16 adults with one run each (16 total) containing 260 volumes and from 19 infant sessions with 1–6 runs each (64 total) containing 6–335 volumes (*M*=112.3). We compared overall SFNR across groups and examined how SFNR changed along the posterior-anterior axis of the brain, which tracks increased distance from the bottom coil (Figure 3).

**Figure 3:**
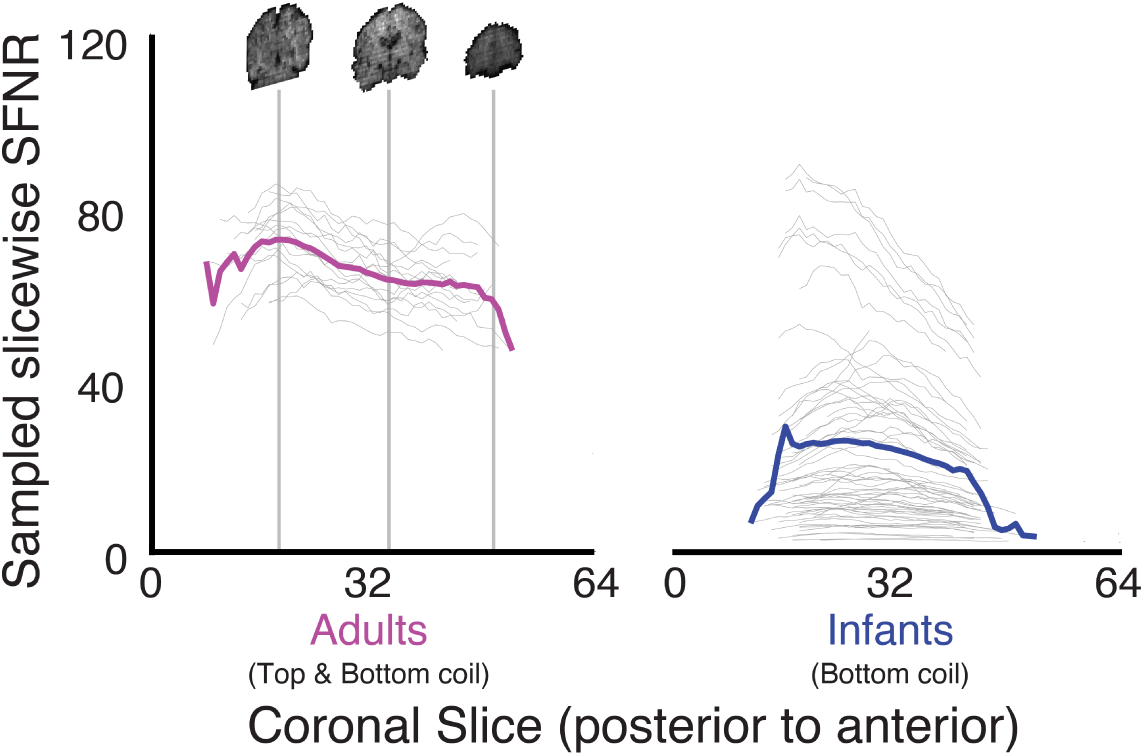
Comparison of SFNR in adults (top and bottom coils) and infants (bottom coil) at different positions along the posterior-anterior axis of the brain volume. SFNR was computed for each voxel and then a random sample of 1,000 voxels from each coronal slice (with at least that many voxels) was averaged. Each gray line is one run from one participant and the colored lines represent the average.

SFNR was higher in adults (*M*=66.0) than infants (*M*=23.6) over the whole brain (*F*(1,77)=76.25, *p*<0.001). This likely reflects factors beyond just the coil, such as age-related differences in head motion and BOLD contrast. Head motion is especially problematic because it can introduce dramatic variability, as a result of signal loss from motion within a volume acquisition and by changing the anatomical location of voxels in the field of view (including across tissue boundaries). Some infants had SFNR in the adult range, suggesting that removing the top coil did not guarantee lower signal quality. Moreover, despite being lower overall, infant SFNR was mostly in a usable range and not dramatically different from the early days of adult fMRI^18^. SFNR was higher in the posterior (*M*=34.2) than anterior (*M*=30.2) half of the brain (*F*(1,77)=36.01, *p*<0.001). If caused by the lack of a top coil, we would expect a larger posterior-to-anterior drop in infants than adults. However, the drop did not differ proportionally (adult *M*=0.90, infant *M*=0.92, Welch’s *t*(35.0)=-0.57, *p*<0.571), and was *smaller* in infants than adults in absolute terms (*F*(1,77)=5.56, *p*=0.021). Thus, we found no additional hit to anterior sensitivity when using only the bottom coil in infants.

### Visual evoked response

We further evaluated data quality by examining basic neural responses. The tasks we tested in these cohorts all involved visual stimuli and so we expected to observe responses in visual cortex. To quantify these responses, we estimated the BOLD activity evoked by task epochs using a general linear model (GLM). To assess the selectivity of visual responses in the brain, we also examined a control region of early auditory cortex (all stimuli were silent).

In Cohort I, there were 14 sessions from 7 unique participants with one or more runs (32 total runs) that contained at least two usable task blocks (*M*=4.5 blocks; range: 2–12). This corresponds to 72.3% of the functional data retained after motion, eye-tracking, and other exclusions during preprocessing. The preprocessed data for each run were trimmed to include blocks of visual stimulation (lasting 36–80s) separated by baseline rest (6s). These task blocks contained a range of stimuli, including faces, shapes, fractals, and objects. Prolonged periods of movie viewing or sleep were not included in this analysis of evoked responses. Blocks of included data were modeled with a canonical hemodynamic response function based on prior infant neuroimaging studies^8,11,20^. Regions of interest (ROIs) were transformed into subject space to measure effects within individual participants, focusing on bilateral early visual cortex (V1), lateral occipital cortex (LOC), and early auditory cortex (A1). To examine group effects across the brain, functional data were aligned to age-specific infant MNI templates, which were in turn registered to adult standard MNI space.

For ROI analyses, we quantified the proportion of voxels in each region that showed a significant visual response within individual runs (Figure 4A). These proportions were reliably different from chance (0.05) across runs in V1 (*M*=0.18 [*SD*=0.22], *p*<0.001) and LOC (*M*=0.14 [*SD*=0.17], *p*=0.001), but not A1 (*M*=0.05 [*SD*=0.09], *p*=0.694). Compared to A1, the proportions were greater in V1 (26/32 runs, *p*<0.001) and LOC (28/32 runs, *p*<0.001); V1 was reliably greater than LOC (22/32 runs, *p*=0.001). Note that requiring reliable responses within individual runs is a conservative test of data quality, as adult data are typically aggregated across multiple runs and participants. Voxelwise analyses examining reliability across runs confirmed these findings and showed that effects were strongest in earlier visual regions (Figure 4B). Similar ROI and voxelwise results were obtained when comparing across entire sessions rather than runs (Figure S2A,B). Despite collapsing across a variety of stimuli and young ages, these findings indicate that our methods can produce reliable visual-evoked BOLD activity.

**Figure 4:**
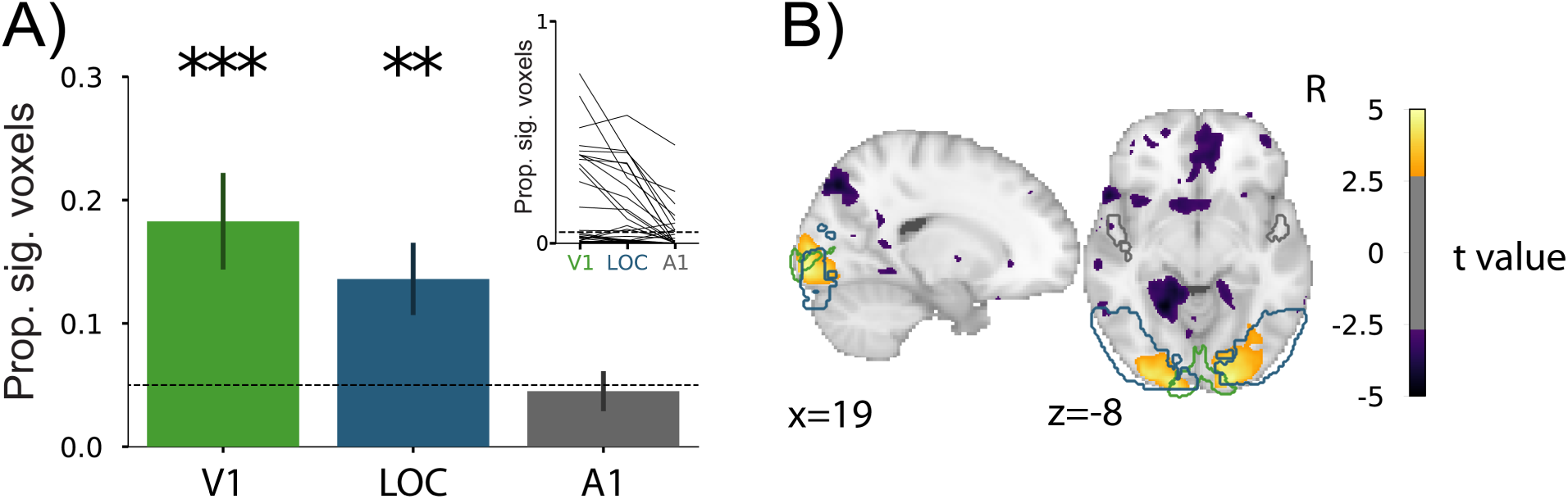
Visual evoked activity for each run from Cohort I. A) Proportion of voxels showing significant visual responses within run (thresholded at *p*<0.05) for V1, LOC, and A1 ROIs (error bars represent standard error across runs). The line plot inset shows the change in proportion of significant voxels across the ROIs for each run. Statistical tests compared against the chance proportion level (0.05): **=*p*<0.01, ***=*p*<0.001. B) Voxels across the whole brain showing reliable responses across runs (*p*<0.005, uncorrected). V1, LOC, and A1 ROIs are outlined in green, blue, and gray, respectively.

### Preprocessing parameters

There is an unavoidable trade-off in infant fMRI between the amount of data retained and the quality of the data included. We chose some parameters to be more liberal than typical adult fMRI (e.g., tolerating motion up to 3mm), knowing that the inclusion of noisy data reduced the power of our analyses. Our reasoning was that infant data are so precious and difficult to obtain that it was worth giving rigorous statistical analyses the opportunity to find signal in the noise. The results described in the preceding sections provide initial evidence that our pipeline could recover signal. Nevertheless, we wanted to explore the impact of our preprocessing decisions more fully. These parameters can be easily modified in the pipeline which makes it possible to explore this trade-off and customize analyses for other tasks and populations. We compared the proportion of significant voxels in the V1 ROI across parameter settings for six potential preprocessing steps using linear mixed models. A higher proportion was not our only consideration: more stringent criteria may improve signal and yet be unacceptable if they exclude too much data. Indeed, longitudinal designs are critical to infant research but would be severely hampered by missing data-points.

The proportion of voxels with significant visual responses varied with the threshold for motion exclusion (Figure 5A, *χ*^2^(4)=9.53, *p*=0.049). Stricter thresholds increased the proportion of significant voxels in V1 (0.5 vs. 3mm: *t*(100.07)=2.06, *p*=0.004; 1 vs. 3mm: *t*(99.77)=2.39, *p*=0.002). However, the amount of data excluded was severe: using 0.5mm as the threshold with the highest proportion, only 14 runs (of 32) across 7 sessions (of 14) were retained with at least two task blocks. This reduced sample size may increase the susceptibility to noise, with a higher proportion of voxels in A1 showing significant visual responses (Figure S3A). The effect of translational motion was limited to the time-points with above-threshold motion themselves. Excluding time-points up to 4s (i.e., 2 TRs) after had no reliable effect (Figure 5B, *χ*^2^(2)=0.12, *p*=0.938), other than reducing the number of time-points retained. Spatial smoothing did affect the proportion of significant voxels in V1 (Figure 5C, *χ*^2^(3)=41.023, *p*<0.001), with our chosen amount reliably better than no smoothing (0 vs. 5mm: *t*(96.00)=-5.19, *p*<0.001). Voxelwise despiking tended to help (Figure 5E, *χ*^2^(1)=6.16, *p*=0.013), while denoising with independent components related to motion hurt (Figure 5D, *χ*^2^(2)=12.57, *p*=0.002; 0.25 vs. 1.0: *t*(96.00)=-3.15, *p*=0.025). Including temporal derivatives in the GLM had a marginal effect (Figure 5F, *χ*^2^(1)=3.78, *p*=0.052). For completeness, we repeated these analyses for LOC (Figure S4) and A1 (Figure S3), and across sessions rather than runs in V1 (Figure S5). Together, these findings indicate that the parameters we chose for our pipeline were sensible, at least with respect to univariate analyses of visual cortex.

**Figure 5:**
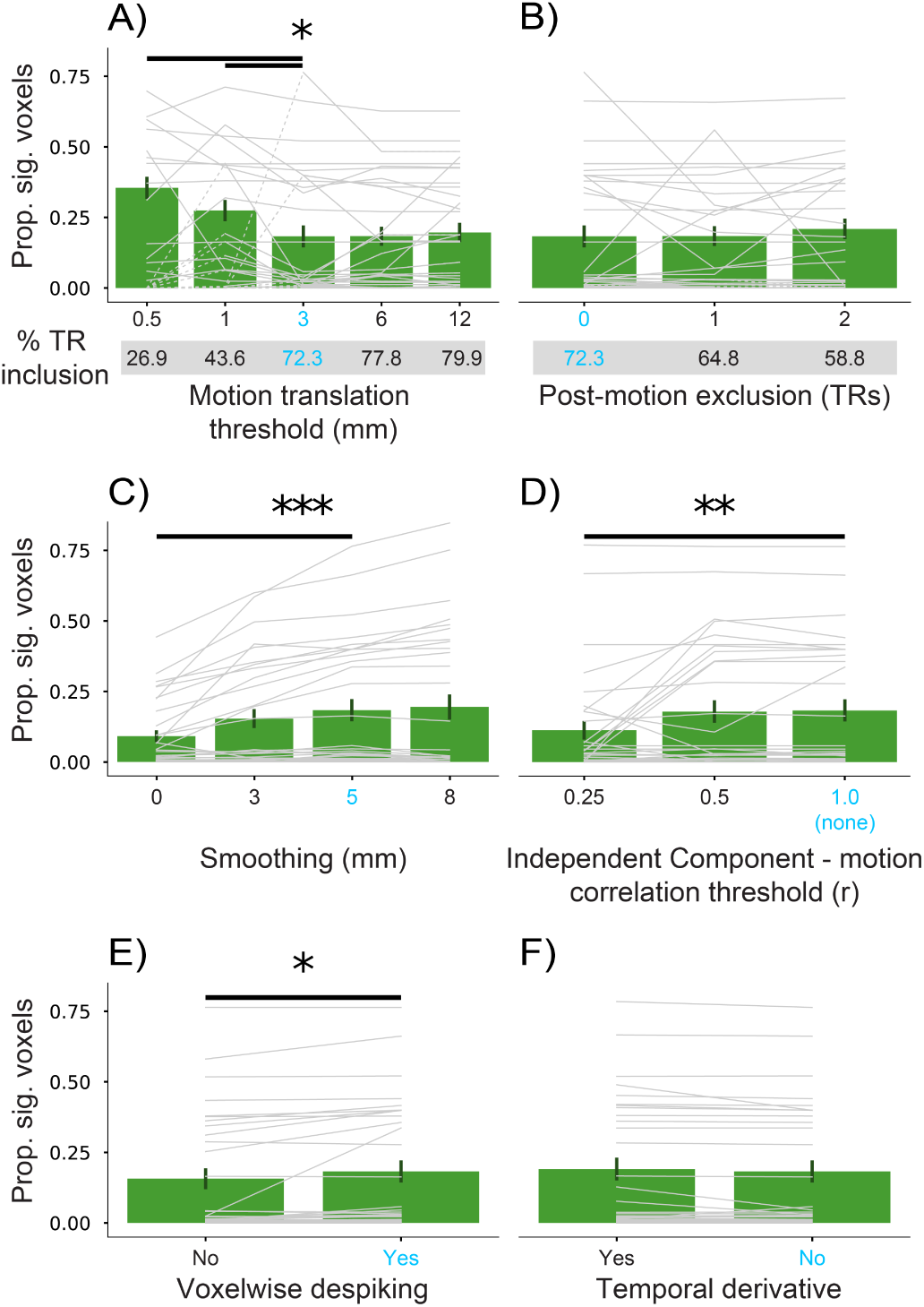
Comparison of the proportion of voxels in V1 showing significant visual responses (*p*<0.05) after various preprocessing decisions. The parameter setting we used for the other analyses is shown in bright blue. Each thin gray line represents a run from one participant. Dashed segments indicate that a run was excluded at that parameter setting (because of insufficient retained data) but included elsewhere. Runs that are excluded entirely regardless of the parameter setting are depicted at zero and not used in the bar plot average. A) Exclusion threshold for time-point to time-point translational motion, and resulting percentage of retained time-points from runs with two or more usable task blocks below. B) Number of time-points into the future removed following above-threshold motion, again with the percentage of retained time-points. C) Full-width half maximum (FWHM) of the Gaussian spatial smoothing kernel. D) Correlation threshold for excluding independent components based on their relationship to motion parameters (lower leads to more components excluded). E) Turning on or off AFNI’s voxelwise despiking. F) Inclusion of temporal derivatives in the design matrix of the GLM. Significance of Chi-square test for omnibus linear mixed model: *=*p*<0.05, **=*p*<0.01, ***=*p*<0.001. Significant differences between our chosen parameter setting (in blue) and other settings indicated by a bold line (*p*<0.05).

### Generalization

Cohort I was collected first and was used to determine preprocessing parameters that balanced data quality and quantity. We then applied these parameters in Cohort II, to test whether they enabled analogous success in new participants, generalizing to a younger population and new site. Cohort II also contained more sessions in which multiple experiments were performed during the same functional run. These runs were trimmed into the time-points corresponding to each experiment and preceding/subsequent rest, leading to experiment-specific *pseudo-runs* that were input to preprocessing. We scanned 15 new individuals between 4 and 10 months old (*M*=6.05 [*SD*=1.8]) across 22 sessions (1–2 sessions per participant). In these sessions, 22.0% of the total time (9.2 mins) produced usable functional and/or anatomical data on average. There was one additional session where the participant did not go into the scanner (resulting in a total of 23 sessions). Of these, 14 sessions resulted in the acquisition of at least one full experiment’s worth of usable data. A variable number of experiments were collected per session (range: 0–3), resulting in a retention rate of 1.04 useful experiment datasets per scheduled session on average.

There were 18 sessions from 13 unique participants with one or more runs (total of 26 runs) that contained at least two task blocks (*M*=4.11 blocks per run; range: 2–12) that could be used to analyze visual evoked responses. The amount of usable data corresponds to 64.6% of the functional data retained after preprocessing with the parameters chosen in Cohort I (blue in Figure 5). We again quantified the proportion of voxels in each bilateral ROI that showed a significant visual response within run (Figure 6A). These proportions were reliably different from chance (0.05) across runs in V1 (*M*=0.12 [*SD*=0.16], *p*=0.008) and LOC (*M*=0.10 [*SD*=0.14], *p*=0.022), but not A1 (*M*=0.05 [*SD*=0.11], *p*=0.975). Compared to A1, the proportions were greater in V1 (14/26 runs, *p*=0.015) and LOC (21/26 runs, *p*=0.031); V1 did not differ from LOC (9/26 runs, *p*=0.390). Voxelwise analyses again revealed reliable neural responses, predominantly in right visual cortex (Figure 6B). Similar ROI and voxelwise results were obtained across entire sessions rather than runs (Figure S2C,D). Thus, we were able to retain a substantial amount of data and recover task-evoked visual responses in young infants with pre-planned acquisition and analysis methods.

**Figure 6:**
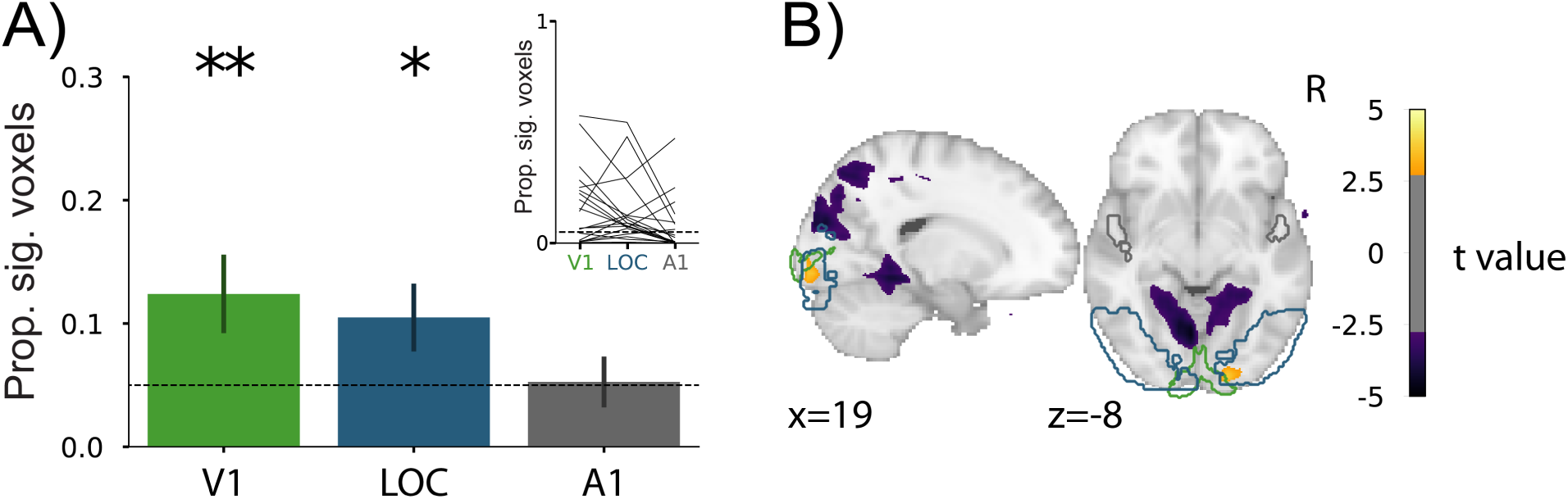
Visual evoked activity for each run from Cohort II (akin to Figure 4 for Cohort I). A) Mean proportion (and standard error) of voxels showing significant visual responses within run (thresholded at *p*<0.05) for V1, LOC, and A1 ROIs. The line plot inset shows the change in proportion of significant voxels across the ROIs for each run. Statistical tests compared against the chance proportion level (0.05): *=*p*<0.05, **=*p*<0.01. B) Voxels across the whole brain showing reliable responses across runs (*p*<0.005, uncorrected). V1, LOC, and A1 ROIs are outlined in green, blue, and gray, respectively.

## Discussion

We presented a suite of novel methods for acquiring and analyzing infant fMRI data, with evidence from two cohorts that these methods can produce high retention rates and reliable neural responses. To address the unique challenges of this population, we adapted the scanning environment and our analysis approach in several ways. Despite deviating from standard protocols and requiring custom equipment and software, these adaptations can be implemented at reasonable cost in most scanning facilities. The main costs include the projector setup, the eye-tracking camera setup, and the vacuum pillow/pump system. Existing projectors and eye-trackers can be re-purposed by adjusting the mounting, calibration, and/or lenses. Indeed, we have implemented these methods successfully at three sites that differed in size and configuration, and we are starting to work with colleagues to implement them at other institutions.

If adopted by the research community, infant fMRI could shed new light on central, long-standing questions about cognitive development by providing more direct access to internal mental representations. This could enable progress in understanding how preverbal infants perceive and categorize the world, comprehend language, predict and simulate, and infer the mental states of others, as well as help explain why as adults we have no memories from this age. To facilitate uptake, we have released our code for data acquisition, a novel and flexible experiment menu system. We have also released our code for data analysis, a semi-automatic pipeline that handles the noise characteristics and unpredictable nature of infant fMRI. These software packages are a key contribution of this work, along with other innovations both technical (e.g., immersive visual display with eye-tracking) and non-technical (e.g., procedures to enhance infant and parent comfort). Although not their original purpose, these tools can also be used in adults for both behavioral and fMRI experiments, including in patient populations. As such, they represent a general-purpose infrastructure for experimental design and analysis both within and outside the field of infant fMRI.

These advances allowed us to acquire considerable fMRI data from awake, behaving infants. We have shared these data, the first substantial public datasets of this type. The two cohorts reflect three years of data collection effort, from 26 unique infants who participated in 45 sessions and completed 57 experiments. The results of individual experiments await further data collection and specialized analysis, and will be reported in forthcoming empirical papers. However, the initial analyses reported here as a proof-of-concept (collapsing across experiment details) revealed reliable and selective univariate visual responses in human infants as young as four months.

Whether our approach is appropriate for other types of analysis (e.g., multivariate), tasks (e.g., auditory), and brain regions (e.g., prefrontal cortex) remains an open question. For example, other groups working with infant fMRI data have used stricter thresholds for motion exclusion^11,12^. Similarly, large motion transients can be excluded at the time-point level, as done here, or used to cleave runs into pseudo-runs with low motion^12^. We hope that the proposed methods, including shared code and data, will increase the feasibility and prevalence of infant fMRI, engaging more researchers to help converge on standards and best practices.

Although these methods proved successful for collecting functional data from awake infants, a limitation of our approach is that anatomical images are generally lower quality than what we obtain from adults or sleeping infants. Given the limited attention span of infants, we prioritized speed over quality when setting up our anatomical sequence. Longer scans that produce higher-quality images in a participant who remains still would, in our experience, result in worse images in infants than shorter scans, because of an increasing likelihood of motion as the scan progresses. The anatomical images we collected were almost always sufficient for aligning functional data, but there is a need for better short anatomical sequences to enable the reconstruction of cortical surfaces and segmentation of subcortical structures. The anatomical scans included in the fMRI datasets we have shared may allow others to improve registration algorithms for images with motion noise. Indeed, this release of infant anatomical scans itself represents a significant increase in the amount of such data publicly available.

In sum, the methods described in this paper, along with the code and data released as companion resources, should make it easier for groups interested in infant fMRI to enter this field. Given our limited understanding of the infant brain, and the success of fMRI in adults, infant fMRI has the potential to provide revolutionary insights into the origins and nature of the human mind.

## Online Methods

### Participants

New data were collected in two cohorts of infants: Cohort I from the Scully Center for the Neuroscience of Mind and Behavior at Princeton University and Cohort II from the Magnetic Resonance Research Center at Yale University. In Cohort I, 11 unique infants aged 6 to 33 months were scanned across 23 sessions (1–8 sessions per participant). Not included in this total were five sessions without fMRI data because the infant would not lie down (4 additional unique infants, 1 infant included above who contributed a usable session on another occasion). In Cohort II, 15 unique infants aged 4 to 10 months were scanned across 22 sessions (1–2 sessions per participant). One other session was excluded because the infant would not lie down (1 additional unique infant). The parent(s) and/or guardian(s) of each infant in a cohort provided informed consent to a protocol approved by the Institutional Review Board of that university. Previously published data^19^ collected from 16 adults at Princeton University were re-analyzed for SFNR comparison.

### Orientation session

When a family was contacted and expressed interest in participating, we first brought them in for an orientation session. This involved meeting a member of our team to discuss research goals, review procedures and safety measures, answer questions, and complete forms (informed consent and preliminary metal screening).

Afterward, we typically introduced the family to the scanning environment using a mock scanner, which consisted of a plastic shell that looked like a scanner but lacked the magnet and other hardware (Psychology Software Tools). The parent placed the infant on their back on a motorized scanner table, which we then slid into the simulated bore. This helped us judge the infant’s ability to lie still. This also helped us judge the parent’s comfort level with separation, though the infant always remained within arm’s reach. We have now shifted away from the mock scanner to a simpler, less cumbersome system that we found to be equally effective in infants. Specifically, we created a simulated bore from a 55-gallon white plastic barrel that was sawed in half lengthwise (into a half circle tunnel), into which we inserted a clear plastic window to show a screen. The parent places the infant on a changing mat inside the tunnel.

We did not use the response to the mock scanner/tunnel for screening purposes. Although we monitored infant behavior and parental comfort throughout, these casual observations were not predictive of future scanning success. Mock scanning can be helpful in children older than two years^21,22^, although not always^23^. Indeed, our experience with infants has been that success is primarily determined by factors that are variable from session to session. This is likely because of dramatic developmental changes every few weeks and because of idiosyncratic factors related to sleep, hunger, illness, teething, time of day, etc.

### Hearing protection

The final part of the orientation, which parents were encouraged to continue practicing at home, was to familiarize the infant with hearing protection. Our goal was to reduce the sound level they experienced during scanning to the range of daily experiences (e.g., musical toys, daycare environments, walking on the street). Sounds around 70 dB are thought to be safe, whereas sounds at or above 85 dB (roughly a loud school environment^24^) could cause damage after extended exposure without protection^25^. Although it is theoretically possible for MRI machines to exceed 110 dB (roughly a loud sports stadium^26^), sound insulation and sequence selection can result in lower levels. Indeed, in our scanning environments and for the sequences we used, the measured sound pressure level reached a maximum of 90 dB.

With hearing protection, it is possible to reduce the sound level by 33 dB, bringing even the loudest possible scanner sounds to safe levels for the duration of the scan. To achieve this noise reduction we combined three forms of hearing protection: First, silicone earplugs (Mack’s Pillow Soft Kids Silicone Earplugs) were inserted into the opening of each ear and expanded over the ear canal (they are tacky and so stay in place better than foam plugs). Second, soft foam cups (MiniMuffs, Natus) were placed over the earplugs and attached to the outer ear with hydrogel adhesive. Third, MRI-safe passive circumaural headphones (MRI Pediatric Earmuffs, Magmedix) were placed over the foam cups. These three layers were intended to provide some redundancy, so that if the earplugs became dislodged the headphones would provide adequate protection and vice versa. Whenever there was a break in scanning, we verified that the hearing protection was intact and re-applied if not. As evidence that the final sound level was comfortable, infants regularly fell asleep during scans and did not startle when scans started, whether awake or asleep. We initially piloted other types of hearing protection outside of the scanner, such as stickers over the tragus, or circumaural headphones that were held around the head by an elastic band, but found that they could not be applied securely and were more prone to failure.

### Scan sessions

We scheduled a scan as soon as possible after the orientation. We left it up to the parent to decide when in the day they thought their child would be best able to participate. They tended to choose times after napping and feeding (often in the morning), though work and childcare constraints and scanner availability also influenced scan time.

Upon arrival, we performed extensive metal screening. Every session, parents filled out a metal screening form for themselves and on behalf of their child, which checked for a medical or occupational history of metal in or on their body. After removing clothing, shoes, and jewelry with metal and emptying their pockets, the parent carried the child into a walk-through metal detector. If the detector sounded, often because of a small amount of metal on the parent (e.g., a clasp), one of the experimenters walked the infant through the metal detector. To double check the child, we passed a high-sensitivity metal-detecting wand, able to find small internal or ingested metal, over their front and back (Adams ER300). We additionally asked the parent if they had seen their child eat anything metallic in the past few days and did not proceed if that was a possibility. Finally, we encouraged parents to bring metal-free toys, pacifiers, and blankets into the scanner to comfort the child; we screened all of these items with the wand before taking them in.

The parent(s) and infant entered the scanner room with one or two experimenters. The hearing protection was put on the infant by one of the experimenters while the other experimenter and parent(s) entertained the child. The infant was then placed on the scanner table. The infant’s head was rested on a pillowcase covering a foam pad in the bottom half of a 20-channel Siemens head/neck coil. The headphones formed a somewhat snug fit reducing the amount of lateral head motion possible, but no additional padding or restraint was used around the head. The infant’s body from the neck down was rested on a vacuum pillow filled with soft foam beads (S & S Technology) covered by a sheet. The edges of the vacuum pillow were lifted and loosely wrapped around the infant to form a taco shape, and the air was pumped out of the pillow until it conformed to the body shape. This bodily stabilization prevented the infant from rolling off the table or turning over, while also reducing body motion during scans. On occasion, and with the recommendation of the parent(s), we swaddled young infants in a muslin blanket before placing them on the vacuum pillow. Overall, however, we found that infants tended to move their head and body *less* when snug but not constricted. The infant’s eyes were covered by an experimenter’s hand while the head was isocentered in the scanner with a laser.

Unlike typical fMRI studies, we did not attach the top half of the head coil. This decision was made for several reasons. It would have obscured the view of the infant’s face from outside the bore, limiting the ability of the parents and experimenters to monitor the infant. The top of the coil would have also blocked the line of sight between the infant and the ceiling of the bore, interfering with our eye-tracking camera and preventing infants from seeing the entire screen projected on the bore ceiling. We also worried that covering the infant’s face would induce unnecessary anxiety and that the hard plastic of the top coil presented an injury risk if the child attempted to sit up.

Figure 1A illustrates the configuration of the research team during infant scanning. We found that it was critical for an experimenter with exceptional bedside manner to remain inside the scanner room adjacent to the parent(s). They monitored and supported infant comfort, provided explanations and directions to the parent(s), and communicated with the experimenters in the control room. This experimenter also adjusted and focused the video camera (12M-i camera, MRC Systems) that was attached to the ceiling of the bore in order to get a clear view of the infant’s eyes. The video feed from the camera was streamed to a monitor placed in the window between the control room and scanner room, which further helped the experimenter and parent(s) in the scanner room monitor the infant.

The experimenter in the scanner room communicated with the research team in the control room about which tasks to run, when to start and stop scans, and how data quality was looking. For Cohort I, the control room spoke over an intercom to the experimenter in the scanner room wearing headphones (Slimline, Siemens), who in turn communicated back with the control room using hand signals visible through the window. For Cohort II, a two-way communication system was used, allowing the experimenter in the scanner room to listen to the control room over headphones (OptoActive II, Optoacoustics) and speak to them through a microphone affixed with velcro tape to the front of the scanner (FOMRI III, Optoacoustics).

The two experimenters in the control room operated computers and equipment. One experimenter controlled the Siemens console computer (e.g., setting up sequences, adjusting alignment, monitoring data quality) and was responsible for communicating with the experimenter in the scanner room. The other experimenter controlled another computer running experimental tasks, the eye-tracker, and the vacuum pump.

### Experiment menu

Given the unpredictability of working with infants, we developed an experiment menu software system that provides complete flexibility in running cognitive tasks during fMRI. This system dynamically generated and executed experimental code in Psychtoolbox^27^ for MATLAB (MathWorks). The experimenter could easily navigate to an experiment from a library of tasks and choose a specific starting block (allowing tasks to be interrupted and resumed), or they could review the progress so far in an experiment. The code coordinated all timing information, receiving and organizing triggers from the scanner, and starting and stopping eye-tracker recordings. After each block ended, there was a short delay before the next block started, allowing the experimenter to determine whether to continue to the next block in the same task, switch to a new task, or stop altogether. It was also possible to rapidly switch to a video at any point, which we showed during anatomical scans, to regain infant interest and attention, and for certain experiments.

This integrated and semi-automated framework for experiments and eye-tracking reduced the burden on experimenters and the possibility of manual errors during already complex procedures. The ability to switch between tasks efficiently within a single ecosystem reduced downtime where the infant was lying in the scanner without any task. This not only increased the amount of time during which usable fMRI data could be collected, but also reduced fussing out that was more likely to occur when nothing was on the screen. Although we developed this system for infants, it could also be used for patient testing and other special populations with similar complications.

The experiment menu system can flexibly incorporate a range of cognitive tasks. Any task that can be designed in Psychtoolbox can be ported to this system, regardless of consideration to response inputs, experiment duration, display parameters, or other factors. To help users understand how the system works, we have provided two sample experiments in the software release that interact with the system in different ways and can be easily modified.

### Eye-tracking

Different types of eye-trackers can be integrated with our software architecture (e.g., EyeLink from SR Research, iViewX from SMI). For Cohort I, we used the frame-grabber capabilities of iViewX eye-tracker software to receive and record input from the MRC video camera. This set-up required manually starting and stopping eye-tracking. For Cohort II, the same camera fed a dedicated eye-tracking computer via a frame-grabber (DVI2USB 3.1, Epiphan). This additional computer ran Python code to save every video frame with a time stamp and was connected to the main experiment computer via ethernet to receive messages, start and stop recording, and perform handshakes. These frames were corrected for acquisition lag and manually coded offline by two or more raters.

To facilitate manual gaze coding, the provided software includes a tool to display relevant video frames offline and convert coded responses into a format compatible with the analysis pipeline. The system was designed to make this laborious task more efficient, allowing coders to quit and resume, accelerate their coding speed, and adjust their field of view. Response code options are entirely flexible (e.g., eyes open vs. closed, fixation vs. saccade, gaze left/center/right, etc.) and could even be used for non-eye behavior (e.g., head motion). This tool also computes coding reliability across raters.

### Ceiling projection

We developed a novel stimulus display system for infants. When using a typical rear-projection system for fMRI, the stimulus is projected on a screen at the back of the bore and the screen is viewed on an angled mirror attached to the top of the head coil. As a result, the stimulus usually covers a small part of the visual field and requires a specific vantage point through the mirror. We could not be sure that these displays would grab the attention of infants. At a more basic level, mirrors may confuse or distract infants. Another approach could be to use a goggle system^11^, which guarantees that the infant can see the stimulus. However, it is hard to monitor the infant with such a system and taking it off can be disorienting.

Instead, we projected visual stimuli directly onto the ceiling of the scanner bore over the infant’s face. We mounted a projector (Hyperion, Psychology Software Tools) approximately six feet high on the back wall behind the scanner, tilted downward to project at the back of the bore. A large mirror placed in the back of the bore behind the scanner table at a low angle reflected the image up onto the bore ceiling, as shown in Figure 1B. This provided a high resolution display (1080p) and a wide field of view stimulation (approximately 115 degrees of visual angle). The thrown image suffered from keystone, elliptical, and stretching geometric distortions, as a result of the angled projection, reflection, and curved bore, but these were corrected automatically in software by a preset screen calibration in the experiment menu code. For Cohort I, we projected directly on the white plastic surface of the bore ceiling. For Cohort II, we taped a piece of white paper to the ceiling to hide the plastic grain. We believe that this large and direct display kept the infants engaged and was natural for them to view. It also gave the parent(s) and experimenters a clear view of the child’s face and allowed for seamless video eye-tracking without calibration. The experiment menu code can be used to set-up and calibrate ceiling projection, but is also compatible with other display types, including rear-projection screens or goggles.

### Tasks

The analyses we report on visual responses combined fMRI data from a variety of stimuli used in several ongoing experiments, including blocks of: looming colorful fractals (Cohort I: 11 runs, Cohort II: 14 runs; 14.6° max size), looming toy photographs (Cohort I: 7 runs, Cohort II: 6 runs; 8°), looming face photographs (Cohort I: 5 runs, Cohort II: 0 runs; 8°), and moving shapes (Cohort I: 9 runs, Cohort II: 6 runs; 10–15°).

Each experimental task was designed to be short, entertaining, and modular. Task blocks generally lasted less than 40 s, though sometimes were longer, as in the case of movies. The tasks used visual effects to maintain attention, including fast motion and onsets (e.g., looming), high contrast textures, bright colors, and relevant stimuli (e.g., faces). Our goal for each session was to obtain up to three full experiments worth of data, which we achieved on occasion. However, the tasks were designed and counterbalanced to provide useful data even when incomplete. Infants sometimes found a given task boring and began fussing or moving, and in such circumstances, we adapted by changing to a new experiment (sometimes later returning to the original experiment). We found that fussing out of one task did not predict that the child would fuss out of other tasks, hence being able to switch tasks within participant increased data yield. The menu system automatically handled timing, scheduled rest periods between blocks/tasks, counterbalanced conditions, and tracked stimulus order and novelty.

At some point during the session, typically after at least one attempted experiment, we collected an anatomical scan. This scan was used for registration of functional data and alignment to anatomical templates. Obtaining a high-quality scan was especially difficult because the infant had to remain still for the entire duration of 3.25 mins (whereas for functional scans, discrete motion only impacted a small number of 2-s volumes). If the infant was awake, we did our best to keep them entertained with either a compelling visual task (e.g., fireworks appearing in different parts of the display) or a movie. If asleep, we blanked the screen. We attempted as many anatomical scans as needed to obtain one of sufficient quality (and as time allowed), though often succeeded in one try.

### Fussiness

Our goal was to make the session as fun and as enjoyable as possible but it was inevitable that some infants got fussy. In our experience, this happened most often at one of three stages: when putting the hearing protection on the infant, when first laying the infant down on the scanner table, and/or when the infant became bored with a task. It was rare that other events, such as the scanner starting, triggered unhappiness. In fact, many infants seemed to be soothed by the scanner sounds and vibrations, and some enjoyed the visual displays so much that they fussed only when removed from the scanner. We did find that talking (loudly) to the infant between scans and patting or holding their hands was soothing. Neither the parents nor experimenters climbed into the bore with the infant. We did not encourage this to avoid distracting the infant and inducing motion or potential confounds. Without such distraction, we found that infants on their own (though within arm’s reach) quickly became enraptured by the visual display. We did allow children to use pacifiers, soothers, bottles, or blankets while in the scanner, which generally had a soothing effect. Although the movement of their jaw while sucking on a pacifier could add noise, this noise was less than that from the motion of an unhappy infant and outweighed the negative impact of otherwise collecting much less data in some sessions.

If a fussy infant could not be soothed or attempted to roll over or climb out, or if the parent(s) asked for a break, we took the infant out of the scanner until they were calm again and ready to resume. The parent would often nurse the infant or give them a bottle or snack, and would change their diaper if needed. In some cases, we had to start and stop 3-4 times before the infant became comfortable enough to provide considerable high-quality data. When the infant had completed all planned experiments, had been in the scanner room for an hour, became too fussy, or fell asleep for a long time (after we completed anatomical scans) we ended the session. In addition to monetary compensation for the family’s time and travel, and a board book for the infant, we also printed a 3-D model of the infant’s brain whenever possible (using Ultimaker 2+ to print surface reconstruction from FreeSurfer^28^). We encouraged families to come in for multiple sessions, and many were happy to do so, generally with one month or more between visits.

### Data acquisition

Infant data were acquired on a 3T Siemens Skyra MRI in Cohort I and on a 3T Siemens Prisma MRI in Cohort II. For anatomical scans, we used a T1-weighted PETRA sequence in all participants (TR_1_=3.32ms, TR_2_=2250ms, TE=0.07ms, flip angle=6°, matrix=320×320, slices=320, resolution=0.94mm isotropic, radial lines=30,000). In two young infants, we additionally piloted a T2-weighted SPACE sequence (TR=3200ms, TE=563ms, flip angle=120°, matrix=192×192, slices=176, resolution=1mm isotropic). For functional scans, we used a T2*-weighted gradient-echo EPI sequence in all participants (TR=2s, TE=28ms, flip angle=71°, matrix=64×64, slices=36, resolution=3mm isotropic, interleaved slice acquisition) covering the whole brain. We did not use multi-band slice acceleration (common in contemporary adult fMRI) because of concerns about peripheral nerve stimulation that could not be reported by our preverbal infants. The adult data for SFNR comparison were acquired on the same scanner as Cohort I and with the same functional sequence, except that the top of the head coil was attached.

### Preprocessing

We developed an efficient analysis pipeline for preprocessing infant fMRI data. This software has been released publicly with this paper and pairs particularly well with the experiment menu system described above. The code is modular and easily editable, while also largely unsupervised. Indeed, any conceivable task-based fMRI experiment could in principle be analyzed with this pipeline. Despite considerable variability in scanning sessions across infants (in fact, no two infants have ever had the exact same session), it typically takes less than two hours of an experienced user’s time to run raw infant fMRI data through the pipeline. Figure S1 depicts the overall preprocessing pipeline schematically. To help users learn how this pipeline works, we provide extensive documentation in a step-by-step tutorial included with the pipeline.

The eye-tracking data we collected from the participants was blindly coded offline by 2–4 naive coders based on the task relevant criteria^29^. Tasks that only required fixation (e.g., movie watching) were coded for whether the eyes were on-screen or off-screen/closed. Tasks that involved viewing images on the left and right of the display were coded for the direction of looking. We calculated reliability by comparing the coded responses between raters. The responses across the coders were aggregated by taking the modal response for a time window of five video frames (100 ms). A tie in the rating counts was resolved by using the code from the most recent previous frame that was not a tie.

After the data from the scanner were converted into NIFTI format, we calculated motion parameters for the functional data. The movement behavior of infants was different from adults because it was often punctate, large in magnitude, and unpredictable, rather than slow and drifting. Hence, the best reference volume to use for motion correction might be different in infants and adults. Specifically, rather than use the first, last, or middle time-point, as is typical in adults, we selected the reference as the volume with the minimum average absolute euclidean distance from all other volumes (the centroid volume). We used FSL^30^ tools (5.0.9 predominantly) for calculating frame-to-frame translations and identified time-points that ought to be excluded because of motion greater than our threshold (3mm).

The stimulus and timing information from each task was converted into FSL timing files using a script. Epochs of data (trials, blocks, or runs) were marked for exclusion at this stage if there was excessive motion and/or if the infant’s eyes were closed for more than half of the time-points in the epoch, or if their eyes were closed during a critical part of the epoch for the experiment. Manual exclusions of data were also specified here, such as when the infant moved out of the field of view of the scanner. The anatomical data were preprocessed using AFNI’s homogenization tools, merged with other anatomical data if available, and skull stripped (AFNI’s 3dSkullStrip^31^).

If more than one task was tested within a run, we created pseudo-runs in which the time-points corresponding to the different tasks were extracted and used to create new run data. This happened more often in Cohort II, in part because we realized between cohorts that we obtained more usable data with less downtime when we scanned continuously rather than stopping arbitrarily when infants finished an experiment. Centroid volumes and motion exclusions were recomputed for these pseudo-runs, which were then input to the preprocessing analyses as if they were collected as separate runs.

First-level analyses were performed to preprocess each run. We started from FSL’s FEAT but added modifications to better accommodate infant fMRI data. We discarded three burn-in volumes from the beginning of each run. We interpolated any time-points that were excluded due to motion, so that they did not bias the linear detrending (in later analyses these time-points were again excluded). We performed motion correction using MCFLIRT in FSL, referenced to the centroid volume as described above. The slices in each volume were acquired in an interleaved order, and so we realigned them with slice-time correction. To create the mask of brain and non-brain voxels we calculated SFNR^18^ for each voxel. This produced a bimodal distribution of SFNR values reflecting the signal properties of brain and non-brain voxels. We thresholded the brain voxels based on the trough between these two peaks. The data were spatially smoothed with a Gaussian kernel (5mm FWHM) and linearly detrended in time. AFNI’s despiking algorithm was used to attenuate aberrant time-points within voxels.

We registered each participant’s functional volumes to their anatomical scan using FLIRT in FSL with a normalized mutual information cost function. However, we found that this automatic registration was insufficient for infants. With this as a starting point, we used mrAlign (mrTools, Gardner lab) to perform manual registration (6 degrees of freedom). One functional run from each session was aligned to the anatomical scan and then each additional run was aligned to the anatomically aligned functional data, all in native resolution. This process was repeated as necessary to improve alignment.

As with registration of functional data to anatomical space, a combination of automatic and manual alignment steps (9 degrees of freedom) were usually needed to register the anatomical scan to standard space (using Freeview from FreeSurfer^28^). The standard space for each infant was chosen to be the infant MNI template closest to their age^32^. These infant templates were then aligned to the adult MNI standard (MNI152 1mm), allowing for comparison between infants and adults in a common space (even if the detailed anatomy does not fully correspond).

After preprocessing and registration, the data can be reorganized into individual experiments. This step is often necessary because any given experiment can be spread across multiple runs (e.g., because of breaks or fussiness). Pseudo-runs from the same task (extracted from different runs with multiple tasks) and entire runs in which only that task was tested are concatenated into a single experiment dataset per participant. We check the counterbalancing across task conditions to prevent biases or confounds related to run number or time. For instance, if an experiment had two conditions, an equal number of epochs from each condition would be selected per run. The voxel time-series for the usable epochs are *z*-scored within run prior to concatenation, to eliminate generic run-wise differences in the mean and variance of BOLD activity. The corresponding timing and motion information are also concatenated. These datasets can then be used as inputs for analyses of individual experiments. Note that we did not perform this step for the data reported in this paper, instead analyzing individual runs from multiple visual tasks (or collapsing across these runs to perform session analyses). Nevertheless we described this step in detail because it is implemented in our software pipeline and will generally be useful for future empirical studies that focus on a particular experimental task.

### Analysis

SFNR was calculated from the raw infant and adult fMRI data. For each voxel in the brain mask, the mean activity was divided by the standard deviation of the detrended activity. The detrending was performed with a second-order polynomial to account for low-frequency drift^18^. To quantify posterior-to-anterior changes in signal, SFNR was estimated for each coronal slice. These slices were taken along the y axis of the acquisition slab, and thus were not precisely aligned with the posterior-to-anterior axis in the reference frame of the head or brain. The average SFNR for each coronal slice (with at least 1000 brain voxels) was computed by sampling the SFNR values of 1000 voxels in that slice. We used this subsampling approach to mitigate differences in infant and adult brain size/voxel counts. The coronal slices were then median split into posterior and anterior halves, which served as a within-subject factor in a repeated measures ANOVA of SFNR, along with age group (infants or adults) as a between-subject factor.

The analysis of visual evoked responses included all of the runs/pseudo-runs that contained at least two task blocks, excluding epochs that were not usable because of motion, eye-tracking, or inappropriate data type (e.g., movie watching or resting state). We fit a univariate GLM across the whole brain for runs/pseudo-runs that contained at least two usable blocks, modeling the response with a task regressor convolved with a double-gamma hemodynamic response function as the basis function. Nuisance regressors were specified for motion relative to the centroid volume and for each excluded time-point. The *z*-statistic map for the task regressor in every run was then aligned into adult standard space.

For the ROI analyses, we defined anatomical ROIs for V1, LOC, and A1 using the Harvard-Oxford atlas. For each ROI and run/pseudo-run, we quantified the proportion of voxels with a *z*-score corresponding to *p*<0.05. The significance of these proportions across runs relative to chance (0.05) was calculated with bootstrap resampling^33^. We sampled with replacement the same number of run proportions from each ROI 10,000 times to produce a sampling distibution of the mean (or mean difference between ROIs). The *p*-value corresponded to the proportion of resampling iterations with mean below chance (or below zero for differences between ROIs). For exploratory voxelwise analyses, we used randomise in FSL to compute a *t*-statistic in each voxel and then applied an uncorrected threshold of *p*<0.005.

In the analyses above, runs/pseudo-runs were treated as independent samples, to assess the statistical reliability of visual responses at the run level. However, there was often more than one run per session, allowing us to additionally examine reliability at the session level. Hence we repeated the same analyses above after concatenating all runs/pseudo-runs within session so that there was only one GLM and set of voxelwise *z*-scores per participant (at a particular age). Despite the smaller number of data-points, the results were largely unchanged (Figure S2).

To explore how preprocessing decisions affected these results, we repeated the univariate analyses above while varying several parameters in our pipeline one at a time: the threshold for excluding individual time-points based on head motion; whether to exclude time-points after instances of motion (after a brain moves there is an imbalance in net magnetization of each slice that can take several seconds to correct); the FWHM of the smoothing kernel; whether to exclude components of the data extracted with independent components analysis (ICA, MELODIC in FSL) that correlate with the six motion parameters (we varied the correlation threshold for regressing out components); whether to use voxelwise despiking to remove aberrant data that can result from motion in voxels near the skull or ventricles; and whether to include the temporal derivatives of the regressors in the GLMs, to account for latency differences.

Some of the preprocessing decisions (e.g., motion threshold) affected our time-point exclusion procedure and in turn the amount of data retained. To calculate these retention rates, we excluded time-points because of above-threshold motion, closed eyes, and when the majority of the block was excluded. Thus, time-points that are otherwise usable but part of an unusable block are considered unusable. For reference, 100% would mean that in runs with at least two usable task blocks, all time-points from all participants were usable. Note that these rates do not account for usable data that were excluded from the analysis of visual responses because the corresponding task was unsuitable for estimating evoked responses (e.g., movie watching).

To compare across parameters for each type of preprocessing, we ran a linear mixed model with condition (parameter) as a fixed effect and run (or session) as a random effect. This approach was chosen instead of a repeated measures ANOVA in order to deal fairly with missing data in cases where a run/session was excluded (e.g., because there were no longer two usable blocks). The Wald Chi Square test was used as an omnibus test to determine whether there were any significant differences in the model, and simple effect comparisons were used to test our default parameter setting against the other options.

### Materials

We have released general software packages that can be used flexibly by other researchers to run new infant fMRI studies, including for data acquisition (github.com/ntblab/experiment menu) and data analysis (github.com/ntblab/infant_neuropipe). We have also shared the specific instantiation of the code (github.com/ntblab/infant_neuropipe/tree/methods validation), as well as the anatomical and functional MRI data from both cohorts needed to generate the results reported in this paper (to be shared on Dryad when published).

## Contributions

C.T.E., N.I.C., V.R.B., & N.B.T-B. initially created the protocol. C.T.E., L.J.S., N.I.C., V.R.B., T.S.Y., & N.B.T-B. collected the data. C.T.E., L.J.S., T.S.Y., & N.B.T-B. developed the pipeline. C.T.E., L.J.S., & T.S.Y. performed the analyses. C.T.E., L.J.S., T.S.Y., & N.B.T-B. wrote the initial draft of the manuscript.

## Acknowledgements

Thank you to all of the families who participated. Thank you to N. Malek for comments on an earlier draft. Thank you to K. Armstrong, J. Bu, C. Greenberg, and L. Rait for recruitment, scheduling, and administration, as well as to the entire Yale Baby School team. Thank you to Y. Braverman, H. Faulkner, J. Fel, and J. Wu for help with gaze coding. Thank you to N. DePinto, R. Lee, L. Nystrom, R. Watts, and N. Wilson for technical support. Thank you to J. Turek for helpful discussions. We are grateful for internal funding from the Department of Psychology and Princeton Neuroscience Institute at Princeton University and from the Department of Psychology and Faculty of Arts and Sciences at Yale University. N.T.B. was supported by the Canadian Institute for Advanced Research.

## Supplementary materials

**Figure S1:**
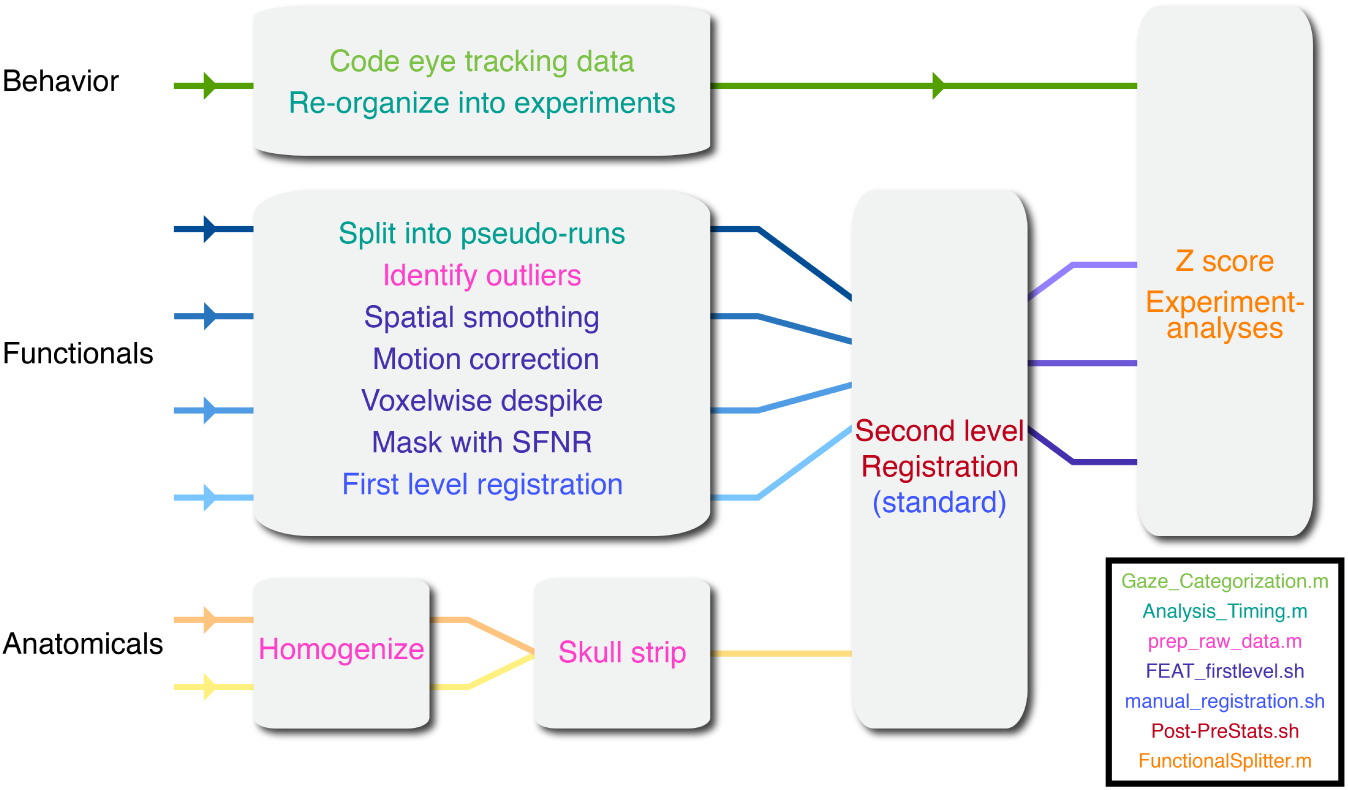
Schematic of the analysis pipeline, color coded by the scripts utilized to perform each operation. This pipeline allows for relatively standardized and automatic processing of infant fMRI data, despite substantial variability across sessions in the amount of data, quality of data, and number and types of tasks.

**Figure S2:**
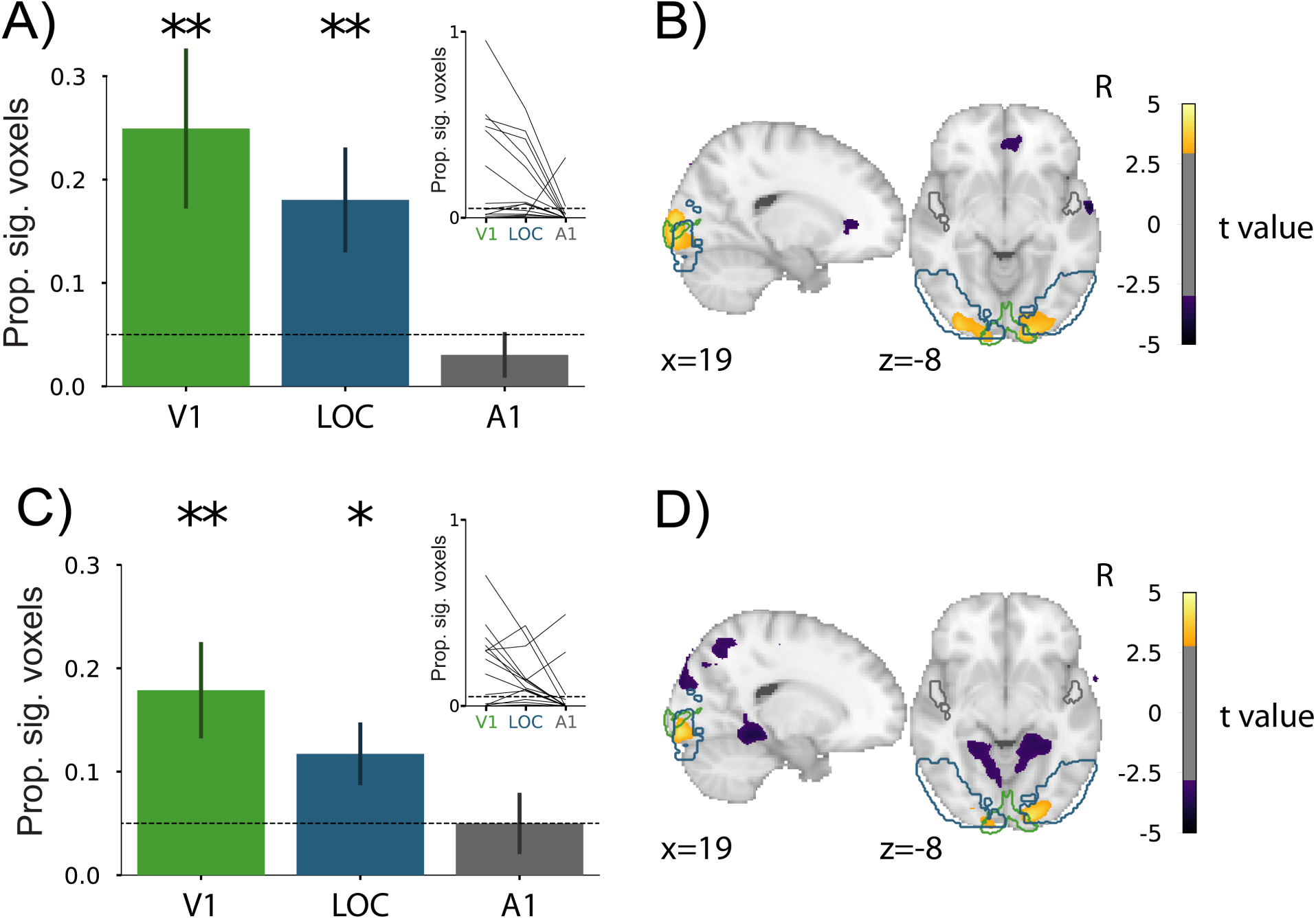
Visual evoked activity for each session from Cohorts I (A, B) and II (C, D). Rather than analyzing each run, here we analyze each session by concatenating the usable blocks from all runs within a session. A) In Cohort I, the proportion of voxels showing significant visual responses within session in V1 (*M*=0.25 [*SD*=0.29], *p*=0.003) and LOC (*M*=0.18 [*SD*=0.19], *p*=0.003) was greater than in A1 (*M*=0.03 [*SD*=0.08], *p*=0.433; V1*>*A1 in 12/14 sessions, *p*=0.005; LOC*>*A1 in 13/14 sessions, *p*=0.009); V1 was also greater than LOC (8/14 sessions, *p*=0.010). The line plot inset shows the change in proportion of significant voxels across the ROIs for each run. B) Voxels across the whole brain showing reliable responses across sessions in Cohort I (*p*<0.005, uncorrected). C) In Cohort II, the proportion of voxels showing significant visual responses within session in V1 (*M*=0.18 [*SD*=0.20], p=0.003) and LOC (*M*=0.12 [*SD*=0.13], p=0.012) was greater than in A1 (*M*=0.05 [*SD*=0.13], p=0.917; V1*>*A1 in 11/18 sessions, *p*=0.002; LOC*>*A1 in 15/18 sessions, *p*=0.027); V1 was also greater than LOC (7/18 sessions, *p*=0.027). D) Voxels across the whole brain showing reliable responses across sessions in Cohort II (*p*<0.005, uncorrected).

**Figure S3:**
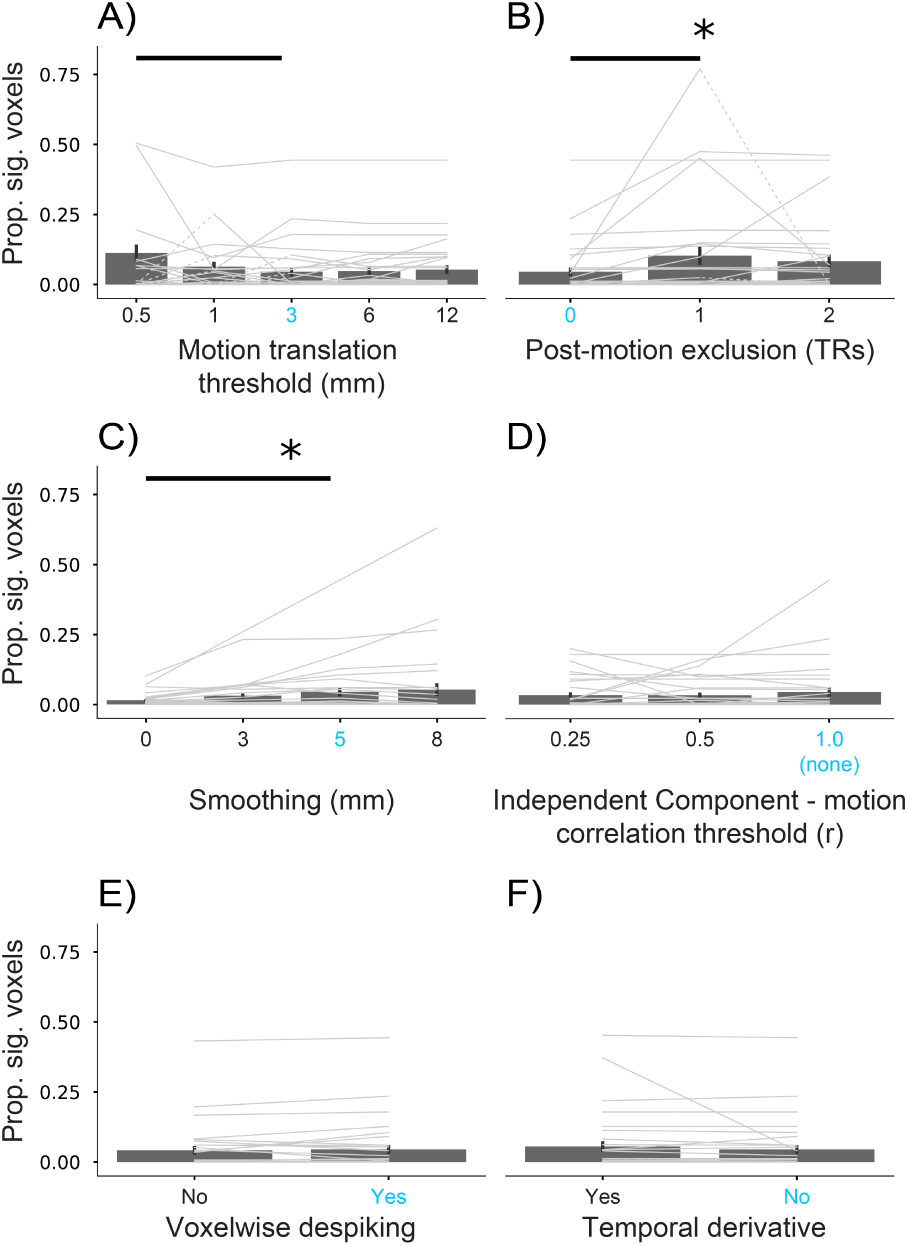
Comparison of the proportion of voxels in A1 showing significant visual responses (*p*<0.05) after various preprocessing decisions. The parameter setting we used for the other analyses is shown in bright blue. Each thin gray line represents a run from one participant. Dashed segments indicate that a run was excluded at that parameter setting but included elsewhere. Runs that are excluded entirely are depicted at zero and not used in the bar plot average. A) Varying the exclusion threshold for time-point to time-point translational motion, with the resulting percentage of retained time-points from runs with two or more usable task blocks (*χ*^2^(4)=4.80, *p*=0.308; 0.5 vs. 3mm: *t*(101.61)=2.05, *p*=0.043). B) Varying how many time-points into the future are removed following above-threshold motion, with the percentage of retained time-points (*χ*^2^(2)=6.60, *p*=0.037; 0 vs. 1: *t*(58.24)=2.48, *p*=0.016). C) Varying the FWHM of the Gaussian spatial smoothing kernel (*χ*^2^(3)=11.30, *p*<0.010; 0 vs. 5mm: *t*(96.00)=-2.48, *p*=0.015). D) Varying the correlation threshold for excluding components from ICA based on their relationship to motion parameters (*χ*^2^(2)=1.23, *p*=0.540). E) Varying whether AFNI’s voxelwise despiking is enabled (*χ*^2^(1)=0.68, *p*=0.408). F) Varying the inclusion of temporal derivatives in the design matrix of the GLM (*χ*^2^(1)=0.91, *p*=0.341). Significance of Chi-square test for omnibus linear mixed model: *=*p*<0.05. Significant differences between our chosen parameter setting (in blue) and other settings indicated by a bold line (*p*<0.05).

**Figure S4:**
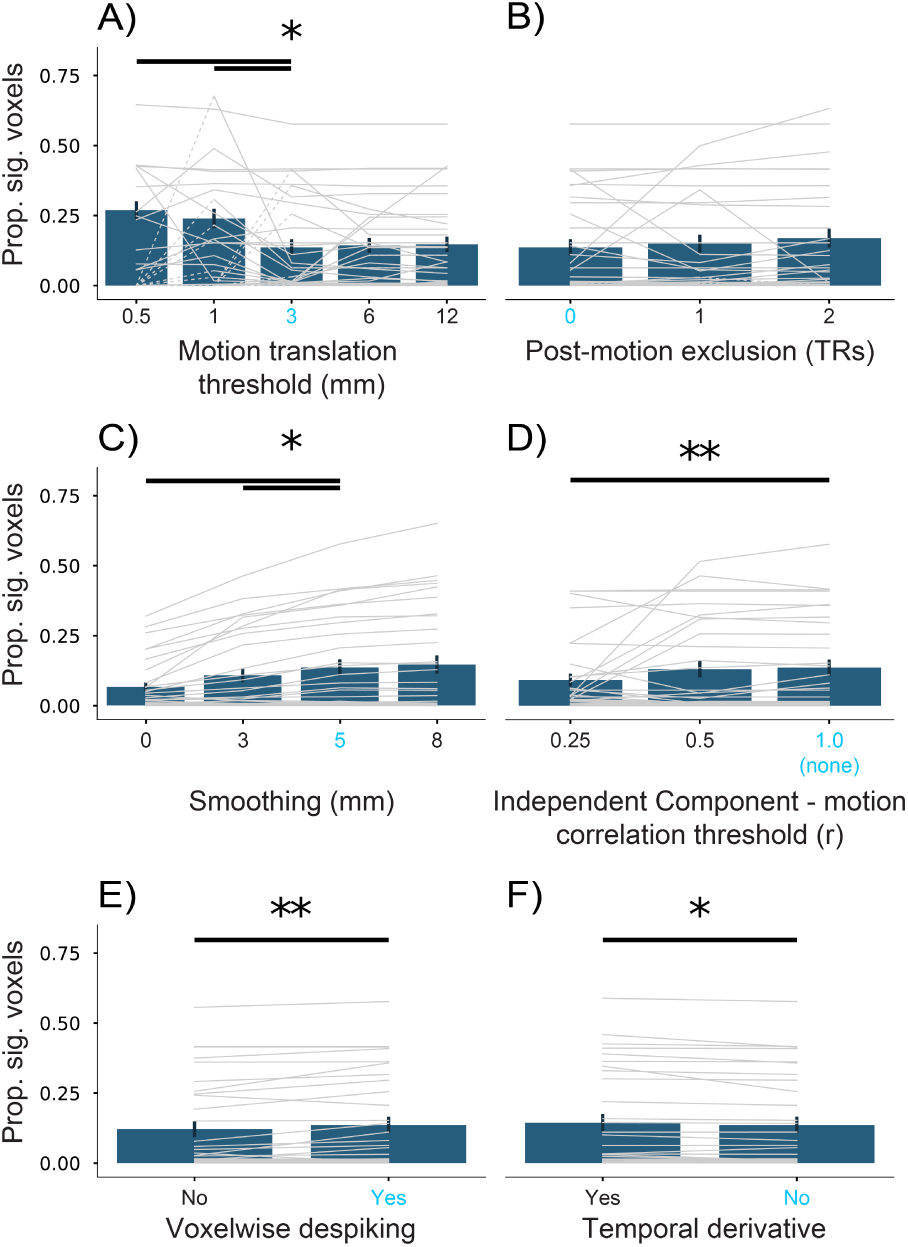
Comparison of the proportion of voxels in LOC showing significant visual responses (*p*<0.05) after various preprocessing decisions. The parameter setting we used for the other analyses is shown in bright blue. Each thin gray line represents a run from one participant. Dashed segments indicate that a run was excluded at that parameter setting but included elsewhere. Runs that are excluded entirely are depicted at zero and not used in the bar plot average. A) Varying the exclusion threshold for time-point to time-point translational motion, with the resulting percentage of retained time-points from runs with two or more usable task blocks (*χ*^2^(4)=12.96, *p*=0.011; 0.5 vs. 3mm: *t*(101.44)=2.09, *p*=0.039; 1 vs. 3mm: *t*(100.85)=3.06, *p*=0.003). B) Varying how many time-points into the future are removed following above-threshold motion, with the percentage of retained time-points (*χ*^2^(2)=0.42, *p*=0.811). C) Varying the FWHM of the Gaussian spatial smoothing kernel (*χ*^2^(3)=51.58, *p*<0.001; 0 vs. 5mm: *t*(96.00)=-5.72, *p*<0.001; 3 vs. 5mm: *t*(96.00)=-2.31, *p*=0.023). D) Varying the correlation threshold for excluding components from ICA based on their relationship to motion parameters (*χ*^2^(2)=7.39, *p*=0.025; 0.25 vs. 1.0: *t*(64.00)=-2.25, *p*=0.015). E) Varying whether AFNI’s voxelwise despiking is enabled (*χ*^2^(1)=8.29, *p*=0.004). F) Varying the inclusion of temporal derivatives in the design matrix of the GLM (*χ*^2^(1)=6.06, *p*=0.014). Significance of Chi-square test for omnibus linear mixed model: *=*p*<0.05, **=*p*<0.01. Significant differences between our chosen parameter setting (in blue) and other settings indicated by a bold line (*p*<0.05).

**Figure S5:**
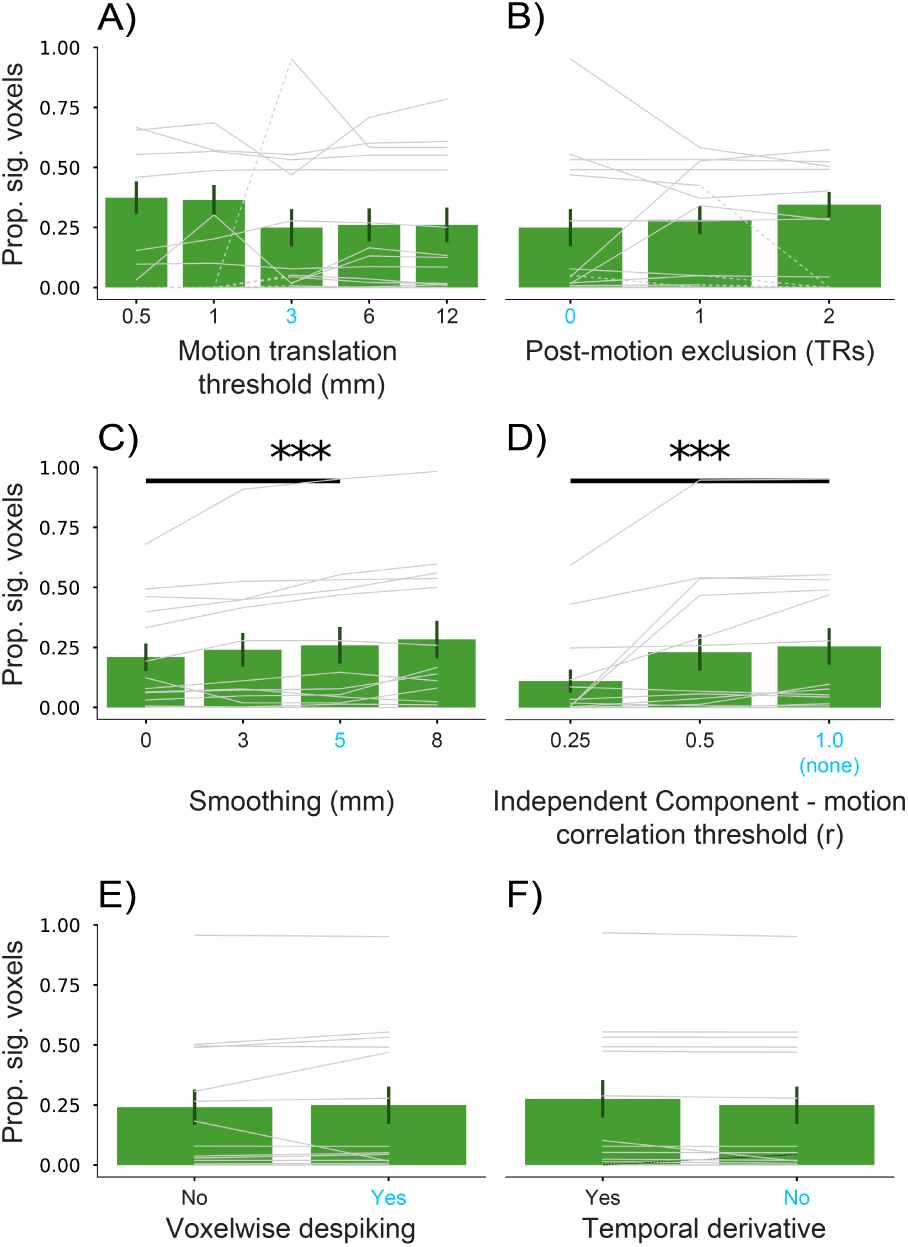
Comparison of the proportion of voxels in V1 showing significant visual responses (*p*<0.05) after various preprocessing decisions for the session-wise analyses. The parameter setting we used for the other analyses is shown in bright blue. Each thin gray line represents a session from one participant. Dashed segments indicate that a session was excluded at that parameter setting but included elsewhere. Sessions that are excluded entirely are depicted at zero and not used in the bar plot average. A) Varying the exclusion threshold for time-point to time-point translational motion, with the resulting percentage of retained time-points from sessions with two or more usable task blocks (*χ*^2^(4)=1.40, *p*=0.845). B) Varying how many time-points into the future are removed following above-threshold motion, with the percentage of retained time-points (*χ*^2^(2)=0.30, *p*=0.861). C) Varying the FWHM of the Gaussian spatial smoothing kernel (*χ*^2^(3)=18.36, *p*<0.001; 0 vs. 5mm: *t*(42.00)=-2.78, *p*=0.008). D) Varying the correlation threshold for excluding components from ICA based on their relationship to motion parameters (*χ*^2^(2)=13.86, *p*<0.001; 0.25 vs. 1.0: *t*(28.00)=-3.48, *p*=0.002). E) Varying whether AFNI’s voxelwise despiking is enabled (*χ*^2^(1)=0.26, *p*=0.612). F) Varying the inclusion of temporal derivatives in the design matrix of the GLM (*χ*^2^(1)=2.78, *p*=0.10). Significance of Chi-square test for omnibus linear mixed model: ***=*p*<0.001. Significant differences between our chosen parameter setting (in blue) and other settings indicated by a bold line (*p*<0.05).

